# Genomic analysis of *Acinetobacter baumannii* prophages reveals remarkable diversity and suggests profound impact on bacterial virulence and fitness

**DOI:** 10.1101/286476

**Authors:** Ana Rita Costa, Rodrigo Monteiro, Joana Azeredo

**Affiliations:** CEB – Centre of Biological Engineering, University of Minho, 4710-057 Braga, Portugal

**Keywords:** Acinetobacter baumannii, bacteriophage, horizontal gene transfer, pathogenicity, antibiotic resistance

## Abstract

The recent nomination by the World Health Organization of *Acinetobacter baumannii* as the number one priority pathogen for the development of new antibiotics is a direct consequence of its fast evolution of pathogenicity, and in particular of multidrug resistance. While the development of new antibiotics is critical, understanding the mechanisms behind the crescent bacterial pathogenicity is equally relevant. Often, resistance and other bacterial virulence elements are contained on highly mobile pieces of DNA that can easily spread to other bacteria. Prophages are one of the mediators of this form of gene transfer, and have been frequently found in bacterial genomes, often offering advantageous features to the host. Here we question the contribution of prophages for the evolution of *A. baumannii* pathogenicity. We found prophages to be notably diverse and widely disseminated in *A. baumannii* genomes. Also remarkably, *A. baumannii* prophages encode for multiple putative virulence factors that may be implicated in the bacterium’s capacity to colonize host niches, evade the host immune system, subsist in unfavorable environments, and tolerate antibiotics. Overall our results point towards a significant contribution of prophages for the dissemination and evolution of pathogenicity in *A. baumannii*, and highlight their clinical relevance.

## Introduction

*Acinetobacter baumannii* was recently indicated by the World Health Organization (WHO) as the number one priority pathogen for research and development of new antibiotics (http://www.who.int/medicines/publications/global-priority-list-antibiotic-resistant-bacteria/en/). This human opportunistic pathogen has been gradually evolving towards clinical success since the 1970s, due to an increasing overall pathogenicity mostly related to a growing multidrug resistance.

Genomically, *A. baumannii* is characterized by a relatively small core genome and a large and diversified accessory genome (1). This indicates gene acquisition and loss as important contributors to *A. baumannii* evolution and adaptation towards pathogenicity. For example, genes associated with antibiotic resistance have been found in both core and accessory genomes of *A. baumannii* (1). In the accessory genome, these genes were found often flanked by integrases, transposases, or insertion sequences, suggesting a possible acquisition by horizontal gene transfer (HGT) from other strains or bacterial species. HGT may thus be a major force in the evolution of *A. baumannii* pathogenicity.

Among mediators of HGT we find bacteriophages (phages), viruses of bacteria thought to be the most abundant biological entities on Earth (2). When infecting a bacterial host, phages may follow a lytic path in which they replicate inside the bacteria and cause cell lysis for progeny release, or a lysogenic cycle where they integrate into the host genome and replicate passively with the bacterial genome. Phages opting for the later are known as temperate phages, or prophages when integrated in the bacterial genome.

Prophages and their bacterial hosts have partly aligned evolutionary interests, since proliferation of the host results in increased prophage population (3). This is possibly the reason why some prophages provide the host bacterium beneficial traits such as protection from infection by other phages (superinfection exclusion) (4, 5), increased host fitness (6), and encoding of virulence factors (VF) exploited for bacterial pathogenesis (e.g. antibiotic tolerance (7) or toxins (8)).

Under certain stimuli, prophages can excise from the host genome, entering the lytic cycle with the release of phage progeny. During excision, a process of specialized transduction may occur, where parts of the bacterial genome adjacent to the prophage may be erroneously excised with the prophage genome and introduced with the virion into a new host (9). Temperate phages thus contribute to host evolution by a constant transfer of genes between host genomes (9, 10). Still, the bacterial host is mindful that prophages can activate their lytic cycle any time, and so prophage genes are under selection for rapid deletion from bacterial genomes. In fact, studies suggest that most prophages in bacterial genomes are to some extent defective (11, 12). Even so, defective prophages can lead to bacterial evolution, with a few bacterial molecular systems thought to derive from the process of prophage inactivation, e.g. gene transfer agents (13), bacteriocins, and type VI secretion systems (T6SS) (14, 15).

The number of prophages in bacterial genomes and their contribution to bacterial evolution differ among species. Here we aimed at evaluating the prevalence of prophages in *A. baumannii* genomes, and at understanding the contribution of these elements to the rapid evolution of pathogenicity in this bacterial pathogen.

## Results

### Prevalence of intact and defective prophages in A. baumannii strains

We analyzed 954 genomes of *A. baumannii* of a total of 1,614 genomes deposited on GenBank at the date of March 2016. Prophages were identified using PHAST and manually curated. A total of 4,860 prophages were found, of which 1,085 were intact and 3,775 were defective. The significantly higher prevalence of defective prophages (Fig. 1A) was expected since intact prophages are usually under strong selection by bacteria for mutations causing prophage inactivation (3). Still, 71.0% of the *A. baumannii* strains contained intact prophages (Fig. 1B), suggesting a recent integration in the bacterial genome.

**Fig 1.**
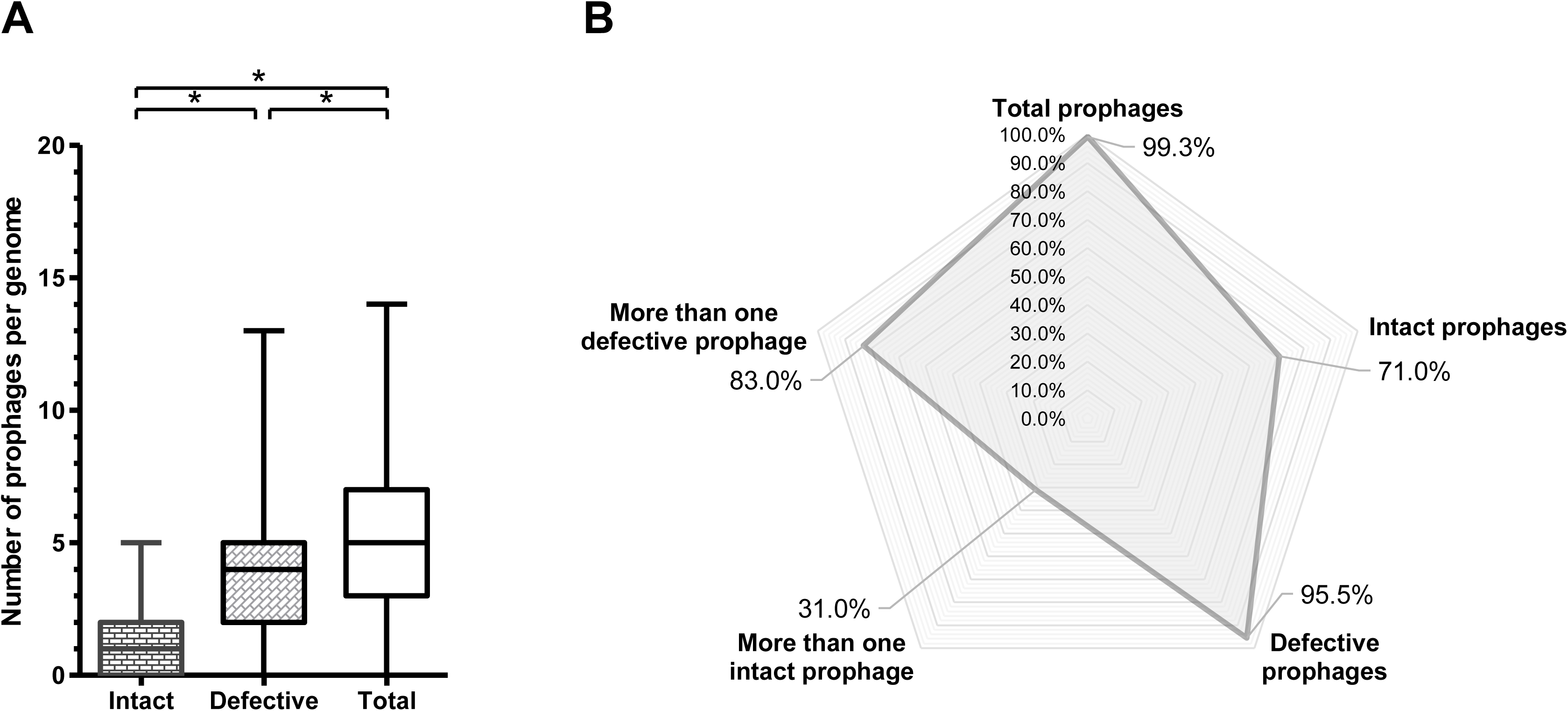
Prevalence of prophages in *Acinetobacter baumannii* genomes. (A) Whiskers plot of prophage frequency per bacterial genome. The horizontal line at the center of the whiskers plot represents the median. The bottom and top of the plot represent the first and third quartiles. The external edges of the whiskers represent the minimum and maximum number of prophages per genome. Significant differences (Tukey’s test) of P < 0.05 are represented by *. (B) Prevalence of total prophages, intact prophages, defective prophages, more than one intact prophage, and more than one defective prophage. Prevalence was determined considering a dataset of 954 *A. baumannii* genomes.

### Distribution of prophages by bacterial genome size

Like Touchon *et al*. (2016) and Bobay *et al*. (2013) did for other bacterial species (16, 17), we question if *A. baumannii* with larger genomes will allow for the integration of more prophages. The existence of more neutral targets for phage integration in larger bacterial genomes may facilitate prophage accumulation without interference with the vital functions of the bacteria (16). To evaluate this hypothesis, we determined the distribution of prophages considering the size of the *A. baumannii* genomes (Fig. 2). A tendency of larger bacteria to harbor increased numbers of prophages can be observed, although the number of prophages appears to stabilize for larger (>4.2 Mbp) genomes (for statistical analysis refer to S1 Table).

**Fig 2.**
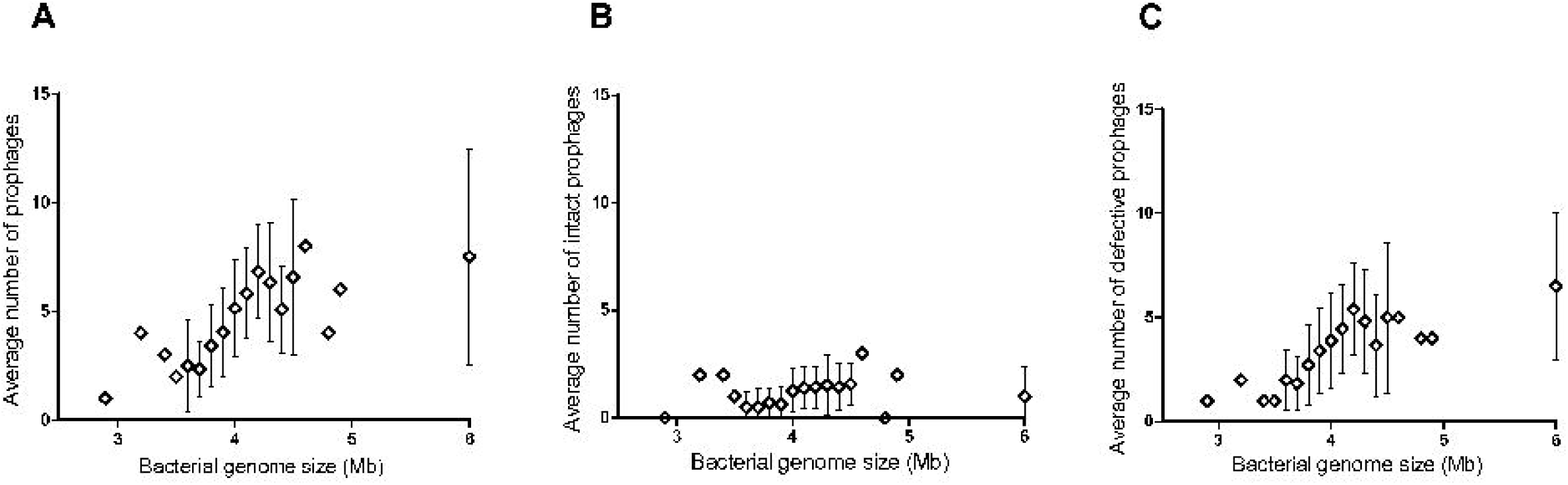
Distribution of prophages among *Acinetobacter baumannii* strains considering the size of the bacterial genome. (A) Average number of prophages (intact and defective) per bacterial genome size. (B) Average number of intact prophages per bacterial genome size. (C) Average number of defective prophages per bacterial genome size.

### Distribution of prophages by family and genome size

We have classed the 138 intact prophages in family taxa based on homology to known classed phages. Classification relied on genes considered the most indicative of family: major capsid protein, large terminase subunit, tail tape measure protein and tail sheath protein. Approximately 66% of the prophages could be assigned a family, with the majority identified as *Siphoviridae*, followed by *Myoviridae* and *Podoviridae* (Fig. 3A). This is in accordance with the estimated distribution in nature (18). On the contrary, the average genome size per prophage family contradicted the literature (http://viralzone.expasy.org/) (Fig. 3B). *Myoviridae* are typically the largest phages, sizing between 33 to 244 kb. However, here *Myoviridae* have the smallest genomes (35 kb, *P* < 0.001) of the prophages with predicted family. Moreover, *Siphoviridae* in *A. baumannii* have the widest size range (32-110 kb) and the largest genomes, when they usually size around 50 kb. Still, in general, the average size of all prophages was 47.6 kb, agreeing with values previously reported for other bacterial species (3, 17, 19).

**Fig 3.**
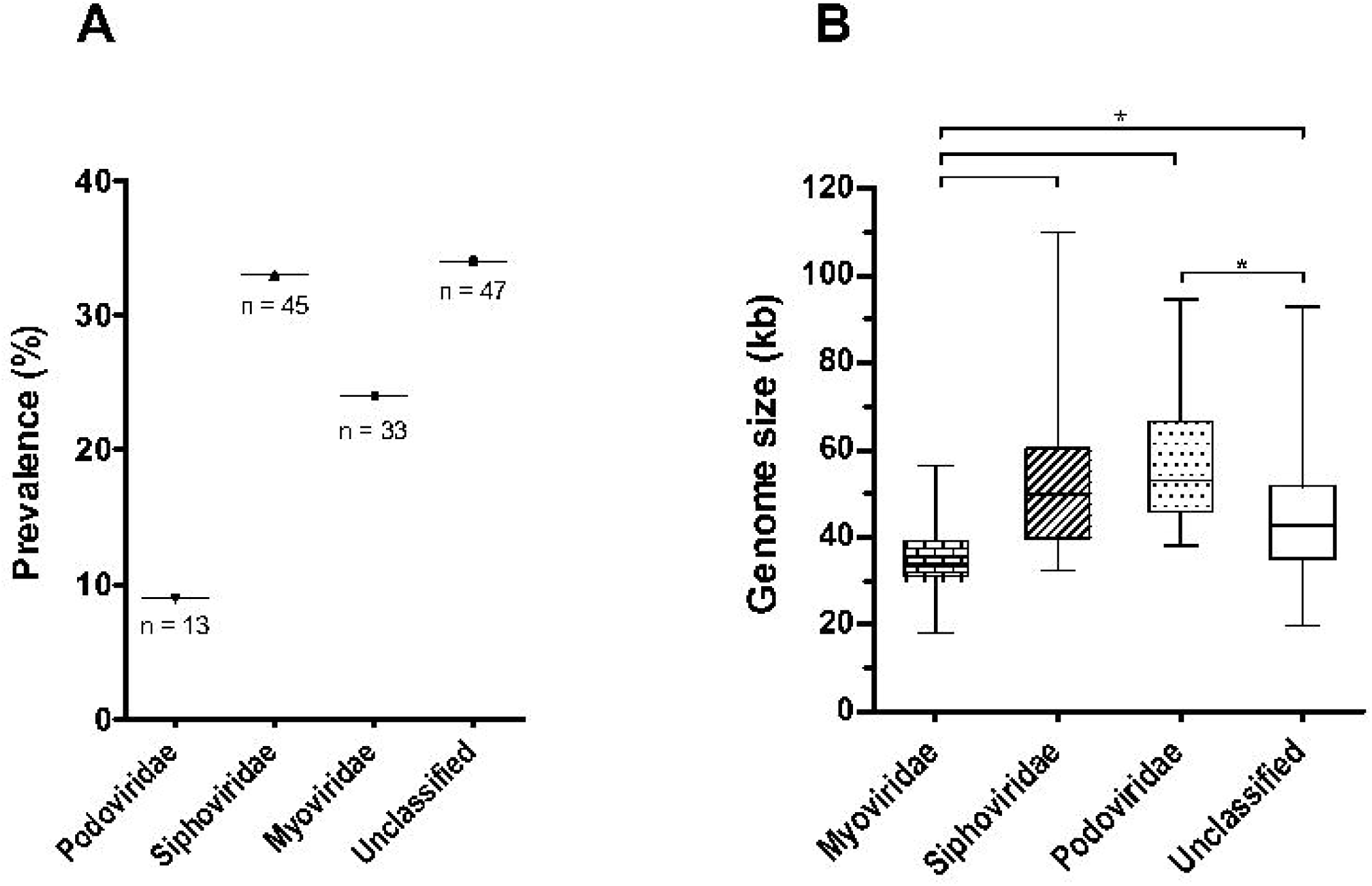
Distribution of genome size of the 138 intact prophages integrating *Acinetobacter baumannii* genomes. (A) Prevalence of prophages in *A. baumannii* genomes by family; (B) Whiskers plot of average genome size of prophages according to family. Horizontal line at the center represents the median, bottom and top of the plot represent the first and third quartiles, and external edges of the whiskers represent the minimum and maximum genome size of prophages per family. Significant differences (Tukey’s test) of P < 0.05 are represented by *.

### Prophages found integrated in A. baumannii mobile genetic elements

We analyzed a set of plasmids associated with the *A. baumannii* strains for the presence of prophages. Four of 74 plasmids were found to possibly harbor intact prophages (S2 Table). Intactness of the prophages is suggested by the presence of proteins related to phage morphogenesis (capsid and tail elements), packaging (terminase), and host lysis (lysozyme), as well as a potential protein involved in phage-host interaction (putative host specificity protein J) (S3 Table and S1 Fig). Curiously, the four plasmids and prophages are highly similar to each other (≥ 82.5% genome homology and ≥ 82.4% proteome homology, see maps in S1 Fig) and to a few other plasmids deposited in GenBank (S3 Table). Some of the *A. baumannii* strains harboring these prophage-containing plasmids have distinct geographical origins (S3 Table), indicating a possible global dissemination of these elements. In fact, since plasmids are much less specific than phages, prophages integrated in these mobile genetic elements may reach a higher diversity of bacteria, and perhaps cross bacterial species.

### Whole genome and proteome comparison of intact prophages reveals remarkable diversity

To determine the relationship and diversity of *A. baumannii* prophages we performed whole genome and proteome dot plot analysis of their sequences. Whole genome analysis revealed 23 small clusters of prophages with genome identity above 50%, indicating strong evolutionary relationships (Fig. 4A). Still, the majority of prophages (about 91% of the comparisons) have genome similarities below 20%, suggesting an enormous genomic diversity. Interestingly, whole proteome analysis revealed an even higher diversity in the amino acid sequences, with about 98% of the comparisons giving a similarity below 10%, and only 9 very small clusters of highly similar prophages (50% identity, Fig. 4B).

**Fig 4.**
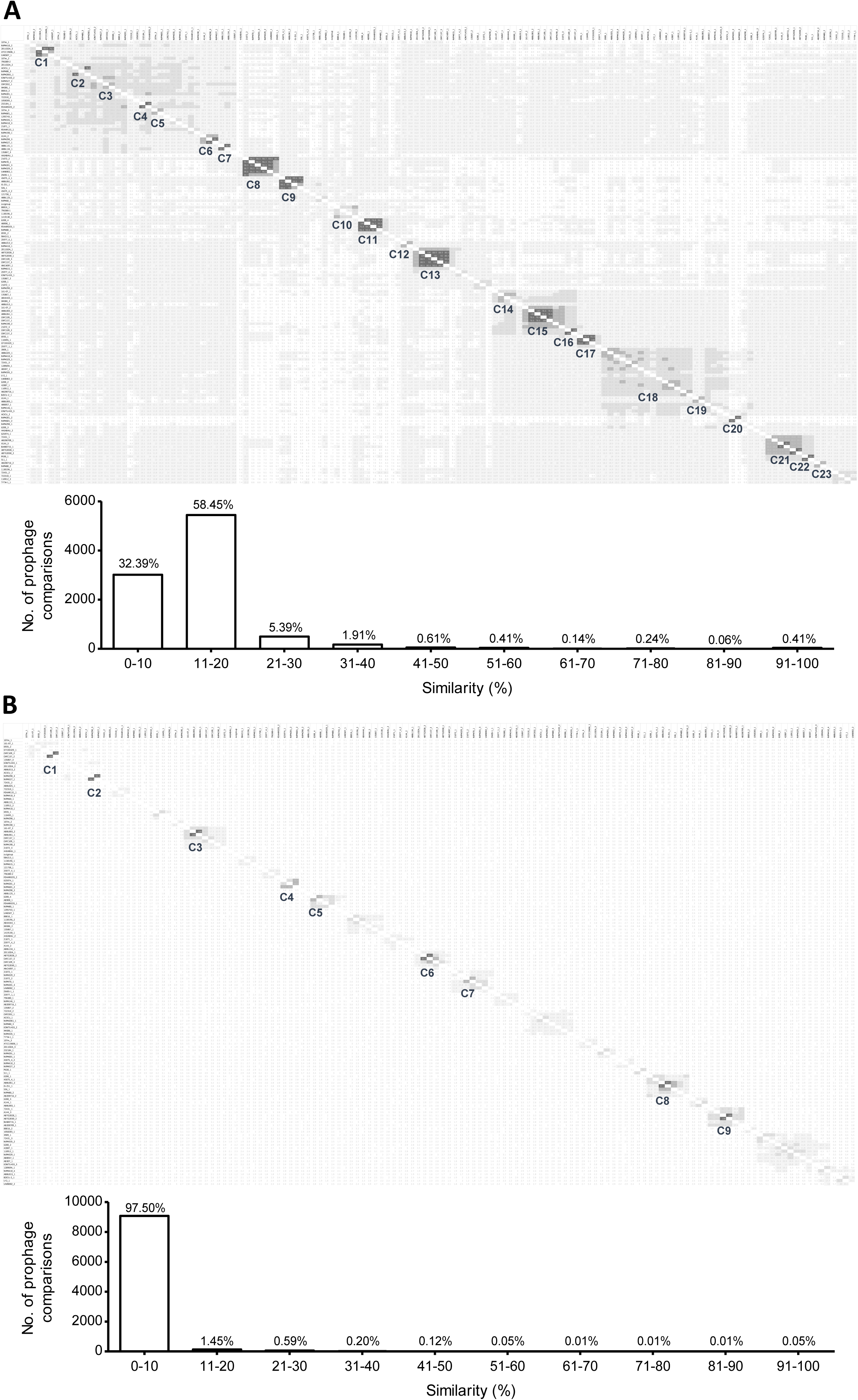
Dot plot matrices of whole sequences of 184 prophages from *Acinetobacter baumannii*. (A) Whole genome analysis; and (B) Whole proteome analysis. Darker zones indicate higher similarity. Clusters of prophages with similarities higher than 50% are indicated and numbered. Graphics summarize the frequency of genome similarity levels found in the analysis. For dot plot matrices with values of similarity see S5 and S6 Tables. Matrices were adapted from the distance matrices retrieved from the phylogenetic trees constructed using Geneious Tree Builder.

To understand if the clusters formed were related to prophage family, we constructed genomic and proteomic phylogenetic trees and inserted family information, as shown in Fig. 5 and Fig. 6. Genomic clusters 1, 4, 6-9, 11, 13, 15-17, 20 and 21 (identified in Fig. 4A) are comprised of sub-clusters containing highly related phages (more than 90% similar). These sub-clusters are identified in Fig. 5 and are composed of prophages of the same family (when determined). Even for areas of lower similarities prophages tend to cluster according to family, although a few singletons are observed. Nevertheless, clusters of the same family are scattered in the tree demonstrating that prophages of the same family can have significantly divergent genomes. A similar analysis is made when observing the proteomics tree (Fig. 6) where sub-clusters of highly related prophages (>90% similarity, clusters 1, 2, 3, 8, and 9) tend to group prophages of the same family. All nine clusters of high proteome similarity are also clusters with high (> 50%) genomic similarity. Moreover, only four of the 13 highly (>90%) similar genome sub-clusters are not identified as clusters in the proteomic analysis. Overall, this demonstrates a strong agreement between both analyses.

**Fig 5.**
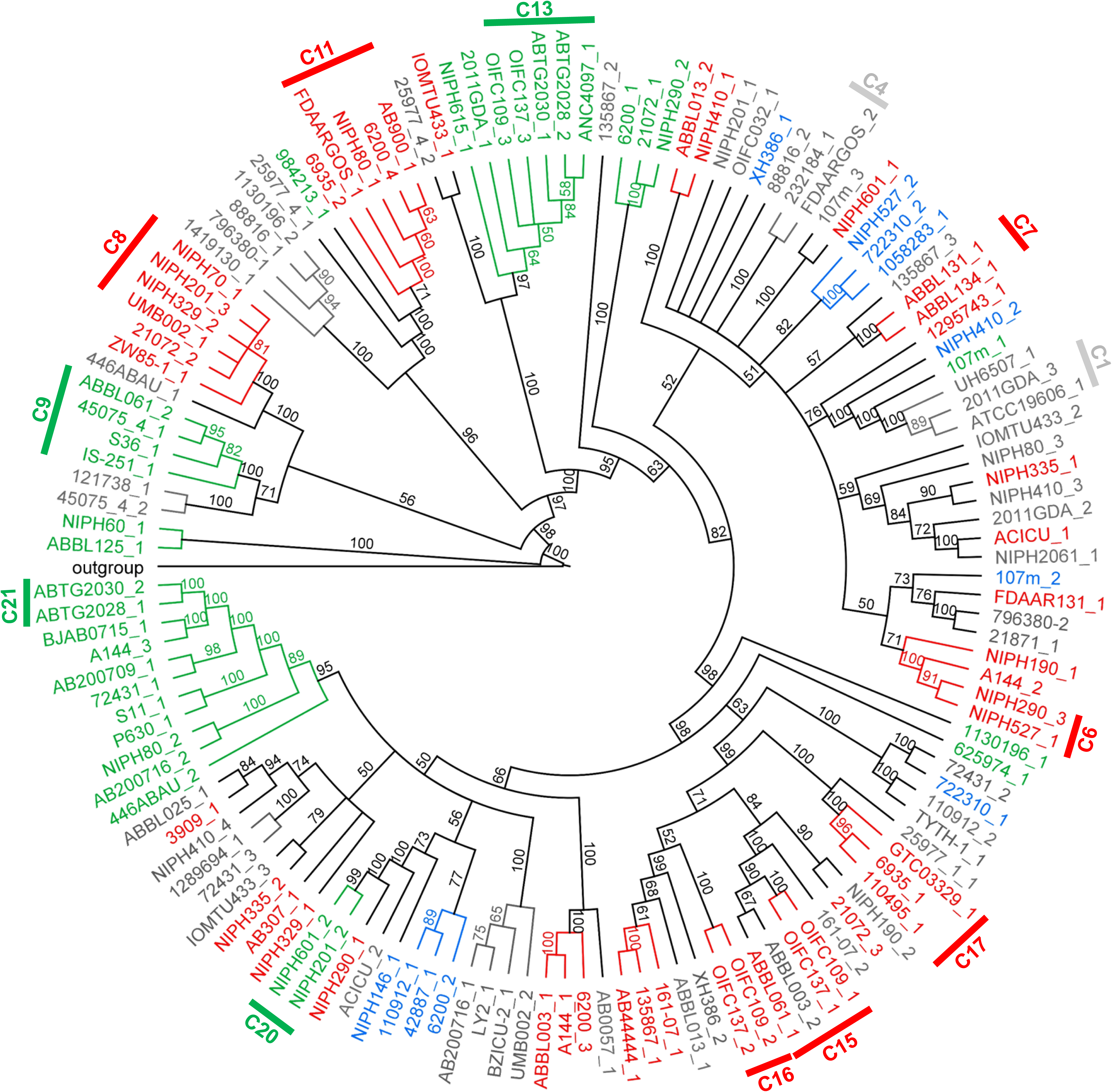
Phylogenetic tree of prophage genomic sequences. Tree was constructed using the Tamura-Nei genetic distance model and the neighbor-joining tree building method in Geneious Tree Builder (Geneious version 9.1.8), with boostrapping set to 100 and tree rooted using *Acinetobacter baumannii* plasmid pNaval18-231 as the outgroup. Branch labels represent bootstrap percentages. Clusters of prophages with genome similarities above 90% are indicated in the tree. Orange: *Siphoviridae*; Green: *Myoviridae*; Blue: *Podoviridae*; Grey: family unknown.

**Fig 6.**
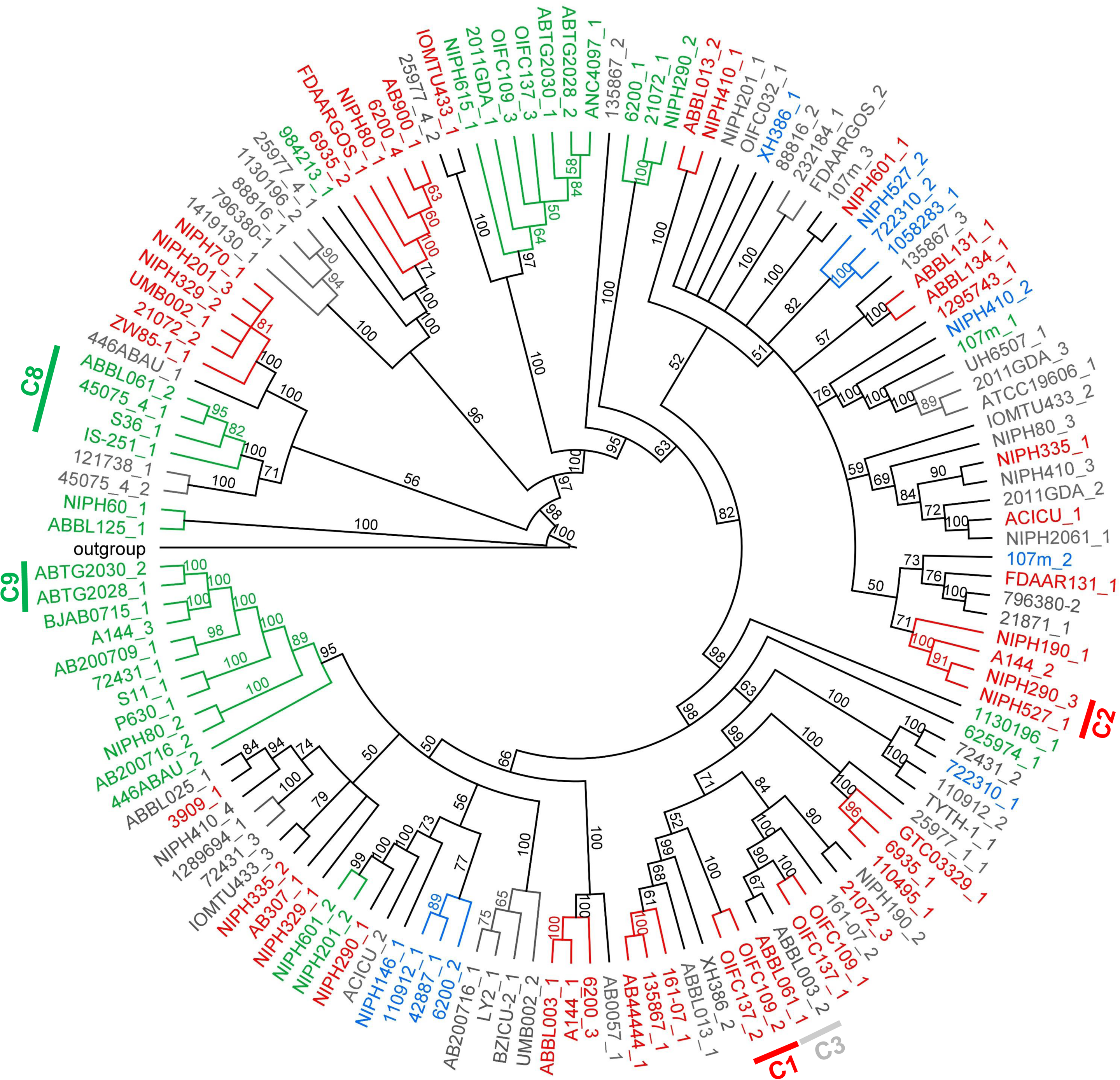
Phylogenetic tree of prophage proteomic sequences. Tree was constructed using the Jukes-Cantor genetic distance model and the neighbor-joining tree building method in Geneious Tree Builder (Geneious version 9.1.8), with boostrapping set to 100 and tree rooted using *A. baumannii* plasmid pNaval18-231 as the outgroup. Branch labels represent bootstrap percentages. Clusters of prophages with proteome similarities above 90% are indicated in the tree. Orange: *Siphoviridae*; Green: *Myoviridae*; Blue: *Podoviridae*; Grey: family unknown.

### Prophages encode a multitude of potential virulence factors

The establishment of stable and long relationships between prophages and the bacterial host has profound implications on both bacterial fitness and virulence (10, 20). Here we hypothesize the rapid spread of pathogenicity in *A. baumannii* to be linked with prophages. We have searched for putative virulence genes encoded by the 138 intact prophages in study. For this purpose, we considered virulence genes as those that might influence bacterial capacity to colonize a niche in the host, evade or inhibit the host immune defense, resist antibiotics, and obtain nutrition from the host; additionally we identified some genes related to fitness. We found that 57% of the *A. baumannii* intact prophages encode putative virulence genes, with an average of 1.6 VF in their genomes (Fig. 7A). By grouping the virulence genes in classes we were able to analyze those most prevalent, as shown in Fig. 7B. A complete list of VF (and fitness factors) identified per prophage can be found at S4 Table.

**Fig 7.**
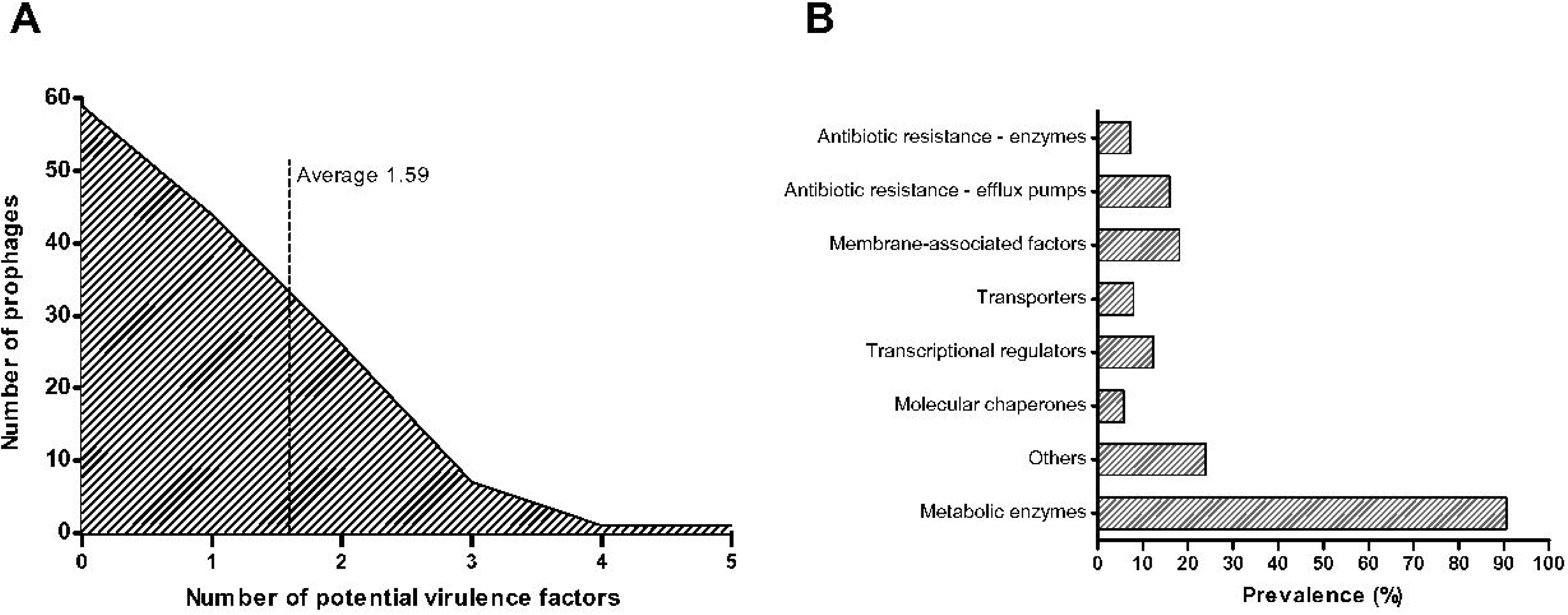
Putative virulence genes identified in the genomic sequences of 184 prophages of *Acinetobacter baumannii*. (A) Distribution of putative virulence genes per prophage; (B) Prevalence of potential virulence factors grouped by class.

The most prevalent putative VF found were membrane-associated factors (18.1%), e.g. outer membrane proteins/adhesins, lipoproteins, fimbrial usher protein, fimbrial biogenesis proteins, and surface antigens. These may interfere with bacterial motility, interaction with host cells and phages, and evasion of host immune defenses (21–23).

Importantly, antibiotic resistance genes were identified in the prophages and can be separated into efflux pumps (15.9%) and enzymes (7.3%). Bacterial efflux systems able to export antibiotics found in the prophages include the major facilitator superfamily (e.g. arabinose efflux permease), the ATP-binding cassette family, the drug/metabolite transporter superfamily, the resistance-nodulation-division family, and the multidrug and toxic compound extrusion family. Additionally, an integral membrane protein TerC, thought to be implicated in the efflux of tellurium ions and thereby in resistance to this antimicrobial (24) was also found in a few prophages.

Resistance to antibiotics also occurs as a result of drug inactivation, drug modification, or target alteration by enzymes (25, 26). Here, the following enzymes were found: beta-lactamase OXA-23 which confers resistance to carbapenems (27); lipid A phosphoethanolamine transferase which provides resistance to cationic antimicrobial peptides (e.g. colistin) (28); acetyltransferase (isoleucine patch superfamily)-like protein which prevents chloramphenicol binding to ribosomes (29); and transcriptional regulators of TetR/AcrR family which regulate the expression of tetracycline efflux and modification proteins involved in antibiotic resistance (30).

Several factors were found that may be involved both in bacterial fitness and virulence. Transcriptional regulators were the most prevalent (12.3%), among which we highlight: LysR family transcription regulators, proposed to regulate a diverse set of genes including those involved in virulence, metabolism, quorum-sensing (QS) and motility (31); IcIR family transcriptional regulators that control genes involved in e.g. multidrug resistance and inactivation of QS signals (32); and bolA, a protein with an important role in the regulation of a network of genes related to biofilm development, flagellar biosynthesis(33), and to the general stress response (33, 34).

Transporters (8.0%) were found with several putative functions, including tolerance/resistance to toxic compounds (e.g. chromate transporter), siderophore export for iron acquisition (TonB-dependent receptor), response to hydroperoxide (vitamin B12 transport periplasmic protein btuE), and interaction with host cells (glutamine transport system permease) (35).

A few molecular chaperones (5.8%) were also identified, with functions mostly related to fimbriae biosynthesis and thus bacterial motility and adhesion to the host (e.g. fimbrial chaperone protein), or response to stress conditions by protecting newly synthesized or stress-denatured polypeptides from misfolding and aggregation (e.g. GroES, GroEL, DnaJ (36)).

A variety of other putative VF were also found, among which proteins involved in red blood cell degradation (e.g. hemolysin activator protein), manipulation of host functions (e.g. Ankyrin repeat protein), promotion of bacterial survival and persistence under stress conditions (e.g. universal stress protein, protein umuD), and targeting of host cells (e.g. RTX toxin).

Additionally, several metabolic enzymes were identified in *A. baumannii* prophages (90.6% prevalence); although not VF they might bring a fitness gain for the survival or proliferation of the host. Among these we highlight: enzymes involved in iron acquisition (e.g. porphyrin biosynthetic protein), which provide an advantage to bacteria in the battle for iron with eukaryotic hosts, especially in nutrient-limited niches (37); enzymes involved in the regulation of bacterial survival under conditions of nutritional (e.g. nucleotide pyrophosphohydrolases (38)) or oxidative stress (e.g. photolyase (39)); enzymes sensing and responding to environmental signals as those resulting from entering the host (e.g. serine/threonine phosphatase (40)); enzymes indirectly involved in antibiotic and xenobiotic resistance (e.g. and acetyltransferase family protein (41)), or in rhamnolipid production, i.e. glycolipids with biosurfactant properties involved in bacterial virulence (anthranilate phosphoribosyltransferase) (42).

## Discussion

As vehicles for HGT, prophages have been linked to bacterial diversification (43) and evolution (20), and may have strong repercussions on bacterial fitness and virulence (9, 10). Only a few studies have characterized the prevalence of prophages in bacterial species and evaluated their role in virulence. Herein we report the analysis of prophage prevalence in *A. baumannii*, and discuss their possible contribute to the evolution of pathogenicity of this human nosocomial pathogen.

We found *A. baumannii* to harbor prophages in most 959 genomes analyzed. While the majority were defective, a high amount of intact prophages were still detected indicating their recent integration. Previous reports (17, 19, 44–46) have estimated lysogen (including intact and defective prophages) prevalence lower than that reported here for *A. baumannii* (99.3%). Still, species as *Streptococcus pyogens* have been reported to have similar high levels of lysogens (90%) (19). It appears that some species are more prone to be lysogenized than others, although the variables associated to the process remain largely unknown. Touchon *et al.* (2016) found minimal doubling time and genome size to be the variables most correlated with lysogeny (17). Fast growing bacteria (with minimal doubling times <2.5 h) were shown to be more lysogenized than slow growing bacteria. Doubling times of *A. baumannii* have been reported to be around 0.5 h (47); as a fast grower a higher percentage of lysogens is therefore expected. An explanation for this phenomenon has been suggested; fast growing bacteria grow weakly under poor environmental conditions (48). In such circumstances phages tend to assume a lysogenic life cycle to preserve their genome while waiting for more propitious conditions for lytic propagation (49). The prevalence of *A. baumannii* in hospital environments, where growth conditions are not ideal, may play a fundamental role in the high prevalence of prophages in *A. baumannii* genomes. Our analysis also suggests that *A. baumannii* strains of larger genomes harbor more prophages, but only for genomes up to 4.2 Mbp. Touchon et al (2016) had similar observations in a set of prophages of mixed species, but stabilization of the number of prophages occurred only above 6 Mbp (17). They have suggested two hypothesis that may also apply here. First, larger genomes may result from selection for functional diversification by HGT, thus facilitating prophage integration. After a certain moment, it is possible that further integration of prophages will not result in the acquisition of novel functions and thus bacteria may become less prone for accepting this type of HGT. Second, superinfection exclusion may be more effective in bacteria with multiple prophages, leading to saturation of prophages in larger genomes. Still, future work is needed to understand the correlation of bacterial genome size and number of integrated prophages.

Among a subset of 138 intact prophages we found *Siphoviridae* to be the most prevalent family, followed by *Myoviridae* and *Podoviridae*, in agreement with the assumed distribution of tailed phages in nature (18). However, the average genome sizes of each prophage family diverged from common descriptions. More strikingly, different trends were observed for each family. *Siphoviridae* had sizes above average and we hypothesize these differences to result from the acquisition of bacterial genes adjacent to the prophage during repeated excision and integration cycles. Conversely, prophages of the *Myoviridae* family have a genome much smaller than the average *Myoviridae* deposited on GenBank. In fact, this family had the smallest average genome size, when it is commonly characterized by the largest phages. A similar observation was made by Bobay et al (2014), although for prophages of different taxa (3). As they have done, we also suggest these phages might have endured some form of genetic degradation that caused a significant reduction of genome size. The reasons why distinct prophage families seem to have evolved differently in the bacterial genomes are unknown. We discarded the hypothesis of inaccurate classifications given by our prophage classification method for two reasons. First, we only attributed prophages with a taxa when comparative analysis of three genes gave concordant classifications. Second, the phylogenetic tree constructed clearly indicates the clustering of phages from the same taxa, supporting our classification. So, further studies are necessary to reveal if this is a common trend among prophages of all bacterial species or if it is specific of *A. baumannii*.

On an interesting note, we found that for a (not so) few prophages, different proteins (e.g. capsid and large terminase proteins) indicated a distinct family, e.g. the prophage of strain ATCC 19606 (759096-823000 bp) identified as *Myoviridae* or *Siphoviridae*, or a prophage of strain NIPH 2061 (1272028-1318975 bp) identified as *Siphoviridae* or *Podoviridae*. We believe this to reflect the mosaic nature of phages, and to suggest the exchange of genetic information by homologous recombination between prophages and infecting phage genomes or other prophages in the same cell. Among other genetic trades, structural genes may be exchanged leading to a difficult interpretation of phage family when exclusively based on genomic information. For example, we found that many prophages having a tail tape measure protein, perceived as characteristic of long tailed phages (50, 51) were classified as *Podoviridae*, e.g. prophages of strains NIPH 146 (3227804-3269564 bp) and NIPH 527 (1517738-1585337 bp). The presence of this gene is therefore non synonymous with long tailed phages, as recently suggested by Ma *et al* (2016) (52)).

Also interestingly, we found some intact prophages integrated in *A. baumannii* plasmids. It is possible that plasmids have acquired the prophages via homologous recombination with the bacterial chromosome, or perhaps by direct integration of the phage into the plasmid. Prophage integration in plasmids may have important implications for the genetic trades occurring within and among bacterial species, resulting in an extremely rich, available gene bank.

Prophages of *A. baumannii* were found to be greatly diversified. Our comparison of 138 intact prophages revealed less than 20% of genome similarity and less than 10% proteome similarity among the majority of prophages. This may indicate one or more of the following: a diversification of prophages into different lineages in ancient times; the constant and intensive diversification of prophage genomes by genetic trades; or a distinct origin of *A. baumannii* prophages (e.g. derived from different bacterial species). Still we could identify a few small clusters of prophages with genomic and proteomic similarities suggesting stronger evolutionary relationships. These may be related to phage taxa, since clusters of high similarity tended to group prophages of the same family.

Some of the genes expressed from prophages can alter the properties of the host, ranging from increased protection against further phage infection, to increased virulence (9). Many cases have been reported linking pathogen virulence to the acquisition of prophages, among which are the well-known *E. coli* O157:H7 whose virulence is correlated with two Shiga-toxin-encoding prophages (53, 54), or *Vibrio cholerae*, producer of the cholera toxin encoded by phage CTXφ (55). Here we found prophages to frequently encode genes of putative function related to bacterial virulence and/or fitness. The prophage-encoded genes may be contributing to the high levels of multidrug resistance found in *A. baumannii*. We identified both drug-specific and multidrug efflux pumps, as well as enzymes able to inactivate/modify the antibiotic or its bacterial target. Among these we highlight the presence of enzymes conferring resistance to colistin, one of the very few last resource antibiotics. Spread of resistance is therefore fostered by prophages, especially under stress conditions that induce prophage excision, as those encountered by bacteria when entering the host environment.

Several other putative VF were found, such as membrane-associated factors, transcriptional regulators, transporters, chaperones, and other proteins, with functions in protection from nutritional and oxidative stress, bacterial motility, interaction with host cells, evasion of host immune defenses, iron acquisition, and regulation of virulence gene expression.

Overall, our results suggest a significant contribution of prophages for the evolution and spread of *A. baumannii* pathogenicity and highlight the clinical relevance of these virions. This study centered the analysis on virulence genes of intact prophages only. However, defective prophages, which are the vast majority, most probably also codify for genes of relevance to *A. baumannii* pathogenicity. Clearly, to understand the evolution of *A. baumannii* as a human nosocomial pathogen we need to recognize and understand the role of prophages in this evolutionary and dynamic process.

## Materials and Methods

### Data collection

A data set of 954 randomly selected complete genomes of *A. baumannii* were retrieved from GenBank (last accessed March 2016). The vast majority of the genomes deposited were at scaffold assembly level, and were used in the analysis as unassembled contigs.

### Detection of prophages in A. baumannii strains

Prophages were detected using PHAST (56). PHAST separates the identified prophages into intact, questionable and incomplete according to criteria that consider the number of coding DNA sequences (CDSs) of a region attributable to prophage CDSs, and the presence of phage-related genes. For the purposes of our analysis, questionable and incomplete prophages were grouped as defective prophages. Furthermore, prophages identified by PHAST were manually curated for increased stringency; only prophages with identified integrase and/or at least one structure gene (e.g. capsid, tail, tail fiber) were considered. Intact prophages were identified as such when all elements required for phage infection were present. Prophages smaller than 10 kb were excluded because these may be difficult to distinguish from other integrative elements (16, 19). The number of intact and defective prophages identified for each strain are detailed in S2 Table.

### Classification of prophages

The prophages of a set of 121 strains were selected for further assessment. Strains were chosen based on the GenBank dendrogram at the date of April 11, 2016 (see S2 Fig). At least one strain from each branch of the dendrogram was chosen. Branch size was considered, i.e. more strains were selected from larger branches. Strains selected harbour a total of 138 intact prophages, whose limits were manually curated using gene annotation and PFAM protein functions. Prophages were classed by comparison of the major capsid protein and large terminal subunit (57–59) to those of previously deposited classed phages, using BLASTp. The presence of elements characteristic of specific families, as the tail sheath of Myoviridae or the tail tape measure protein of long tailed phages(9), was also considered for prophage classification. Prophages were attributed a family only when the results of all comparisons gave identical classifications. Prophage classification can be seen in S4 Table.

### Whole genome and proteome comparisons

Prophage genomic and proteomic sequences were aligned using MAFFT version 7.304 (60), using the Phylip output format, sorted, strategy “—auto”. The genome phylogenetic tree was constructed using the Tamura-Nei genetic distance model and the neighbor-joining tree building method in Geneious Tree Builder (Geneious version 9.1.8 (61)). The proteome phylogenetic tree was constructed using the Jukes-Cantor genetic distance model. Boostrapping was set to 100 and the trees were rooted using *A. baumannii* plasmid pNaval18-131 as the outgroup.

### Identification of potential virulence factors encoded by prophages

The subset of 138 intact prophages was analysed for the encoding of putative VF. For this, prophage proteins with assigned function (by myRAST and sequence comparison using PFAM and HMMER) were correlated to putative VF using PubMed search. A full list of the identified putative VF can be found at S4 Table.

### Statistical analysis

Statistical analysis of the data was performed using the independent samples t-test for comparisons of two samples, or one-way analysis of variance (ANOVA) with post-hoc Tukey HSD test for comparing multiple variables, using the software GraphPad Prism 5, and considering a significance level of 95 %.

## Acknowledgments

This work was supported by the Portuguese Foundation for Science and Technology (FCT) under the scope of the project PTDC/BBB-BSS/6471/2014 (POCI-01-0145-FEDER-016678), the strategic funding of UID/BIO/04469/2013 unit and COMPETE 2020 (POCI-01-0145-FEDER-006684). This work was also supported by BioTecNorte operation (NORTE-01-0145-FEDER-000004) funded by the European Regional Development Fund under the scope of Norte2020 – Programa Operacional Regional do Norte. Ana Rita Costa acknowledges FCT for grant SFRH/BPD/94648/2013. The funders had no role in study design, data collection and analysis, decision to publish, or preparation of the manuscript. The authors declare no competing interests.

## Supporting information

**S1 Fig. Map of plasmids of Acinetobacter baumannii containing prophages**. (A) Plasmid p6200-114.848kb of *A. baumannii* strain 6200; (B) Plasmid pAB386 of *A. baumannii* strain XH386; (C) Plasmid pABTJ2 of *A. baumannii* strain MDR-TJ; (D) Plasmid ZW85p2 of *A. baumannii* strain ZW85-1. Green arrows: identified open reading frames; Yellow arrows: annotated coding DNA sequences (CDS); Pink arrow: tRNA; Red region: prophage sequence identified. Graphic for GC (blue) and AT (green) content in center.

**S2 Fig. Dendogram of *Acinetobacter baumannii* retrieved from GenBank at the date of April 11, 2016.** Representative strains from each branch were selected for detailed analysis of encoded prophages.

Table S1. Significant differences (P < 0.05) on the number of total, intact, or defective prophages harbored by Acinetobacter baumannii strains of different genome sizes.

S2 Table. Number of intact and defective prophages identified for each *Acinetobacter baumannii* strain and respective plasmids.

S3 Table. Features of prophages found in plasmids of *Acinetobacter baumannii*.

S4 Table. Family, virulence and fitness factors identified for 138 prophages of *Acinetobacter baumannii* strains.

S5 Table. Distance matrix of whole genome sequences of 138 prophages from *Acinetobacter baumannii*. Similarity values are presented in percentages, and darker zones indicate higher similarities.

S6 Table. Distance matrix of whole proteome sequences of 138 prophages from *Acinetobacter baumannii*. Similarity values are presented in percentages, and darker zones indicate higher similarities.

