## Supplementary Materials for "Genomic analysis of *Acinetobacter baumannii* prophages reveals remarkable diversity and suggests profound impact on bacterial virulence and fitness"

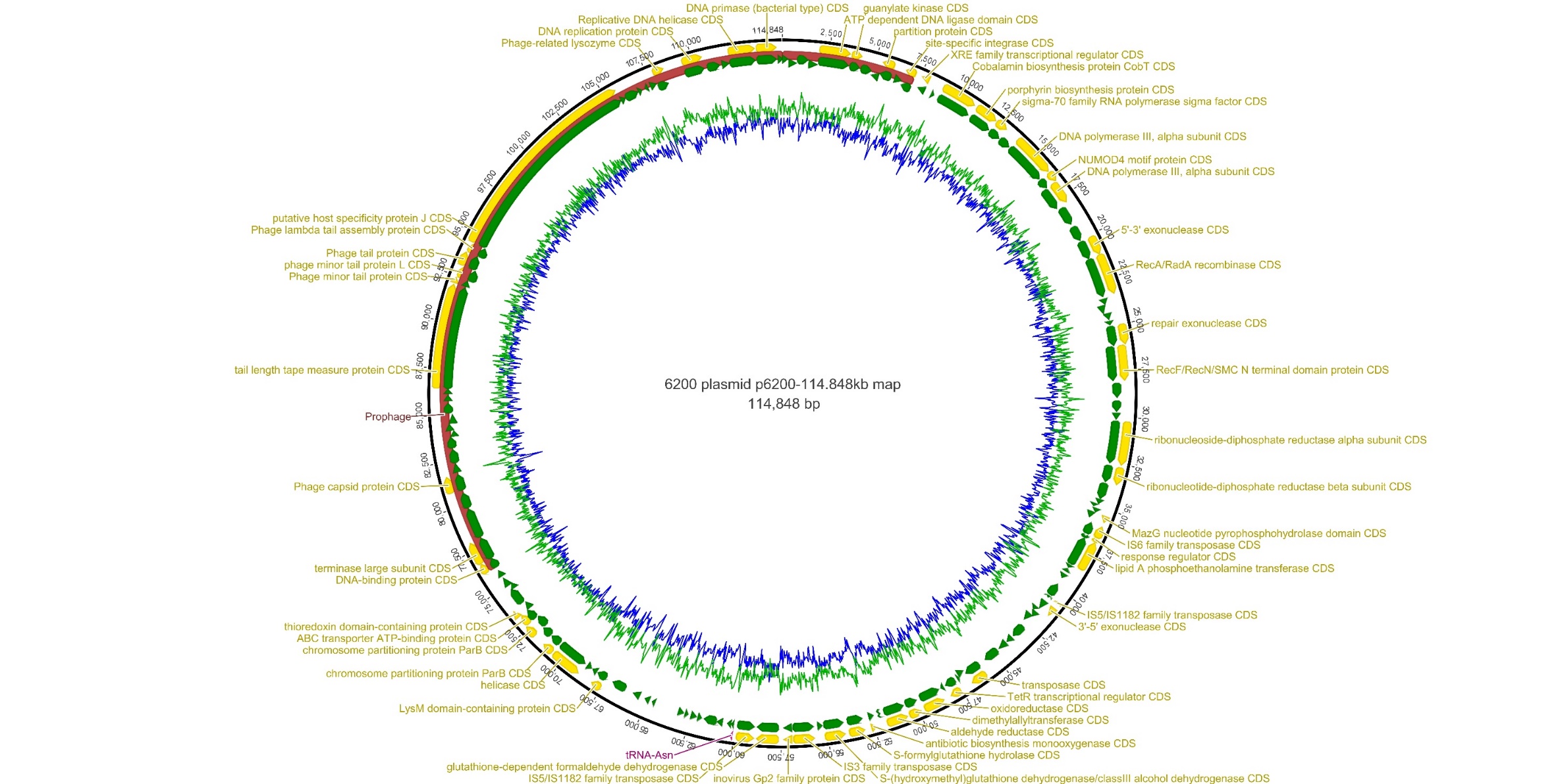

**Figure S1A.** Map of plasmid p6200-114.848kb of *Acinetobacter baumannii* strain 6200. Green arrows: identified open reading frames; Yellow arrows: annotated coding DNA sequences (CDS); Pink arrow: tRNA; Red region: prophage sequence identified. Graphic for GC (blue) and AT (green) content in center.

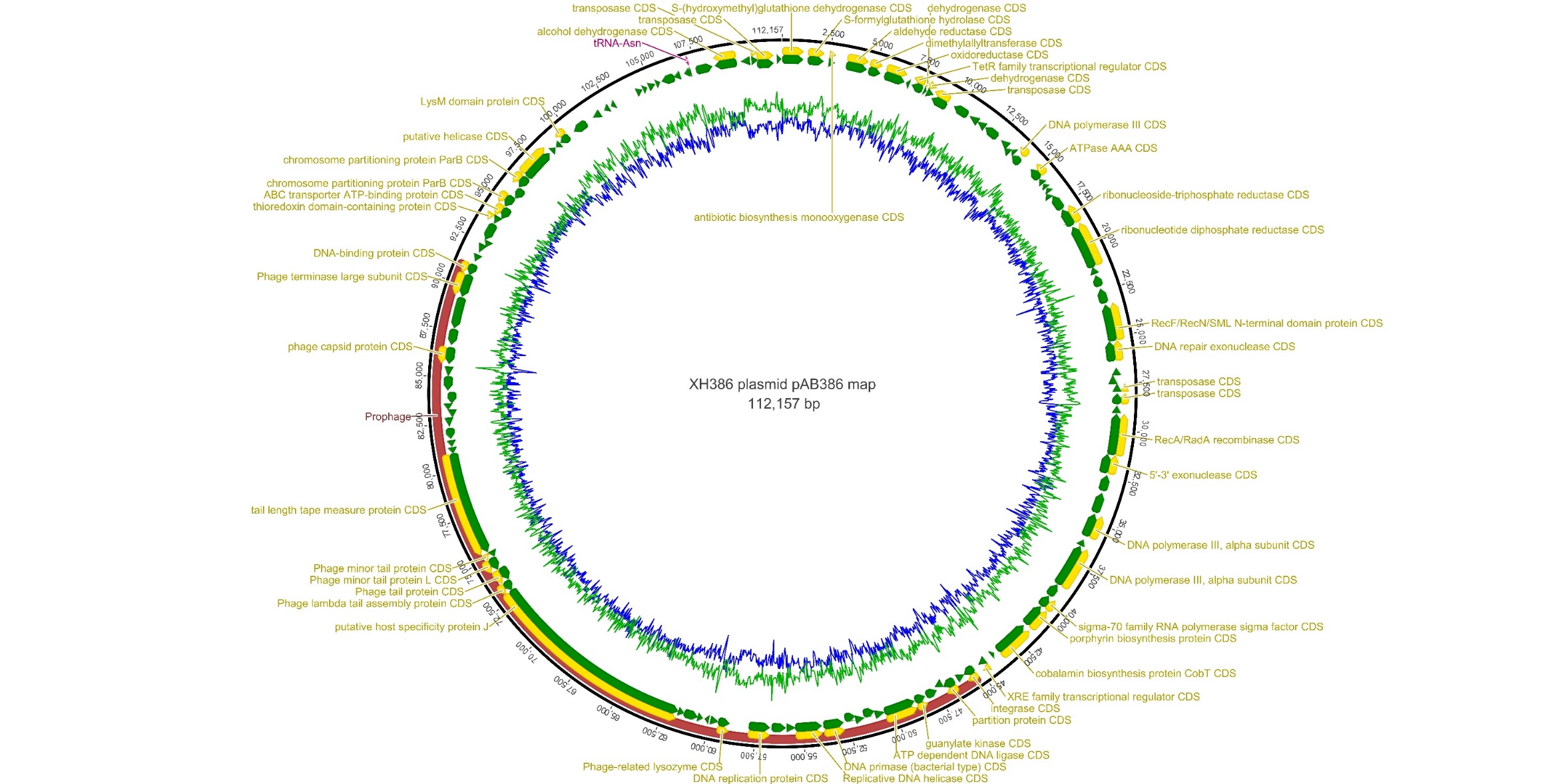

**Figure S1B.** Map of plasmid pAB386 of *Acinetobacter baumannii* strain XH386. Green arrows: identified open reading frames; Yellow arrows: annotated coding DNA sequences (CDS); Pink arrow: tRNA; Red region: prophage sequence identified. Graphic for GC (blue) and AT (green) content in center.

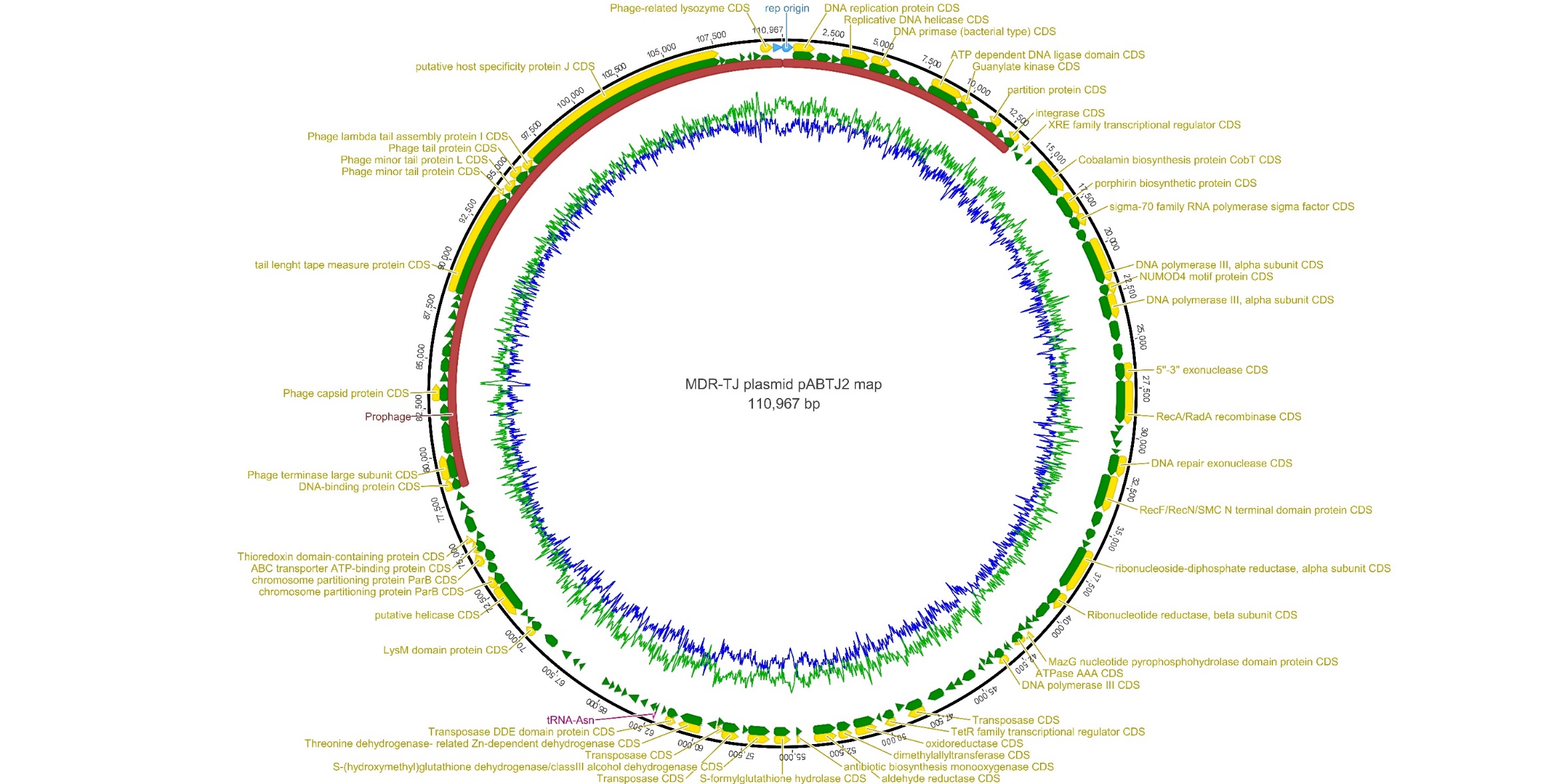

**Figure S1C.** Map of plasmid pABTJ2 of *Acinetobacter baumannii* strain MDR-TJ. Green arrows: identified open reading frames; Yellow arrows: annotated coding DNA sequences (CDS); Pink arrow: tRNA; Red region: prophage sequence identified. Graphic for GC (blue) and AT (green) content in center.

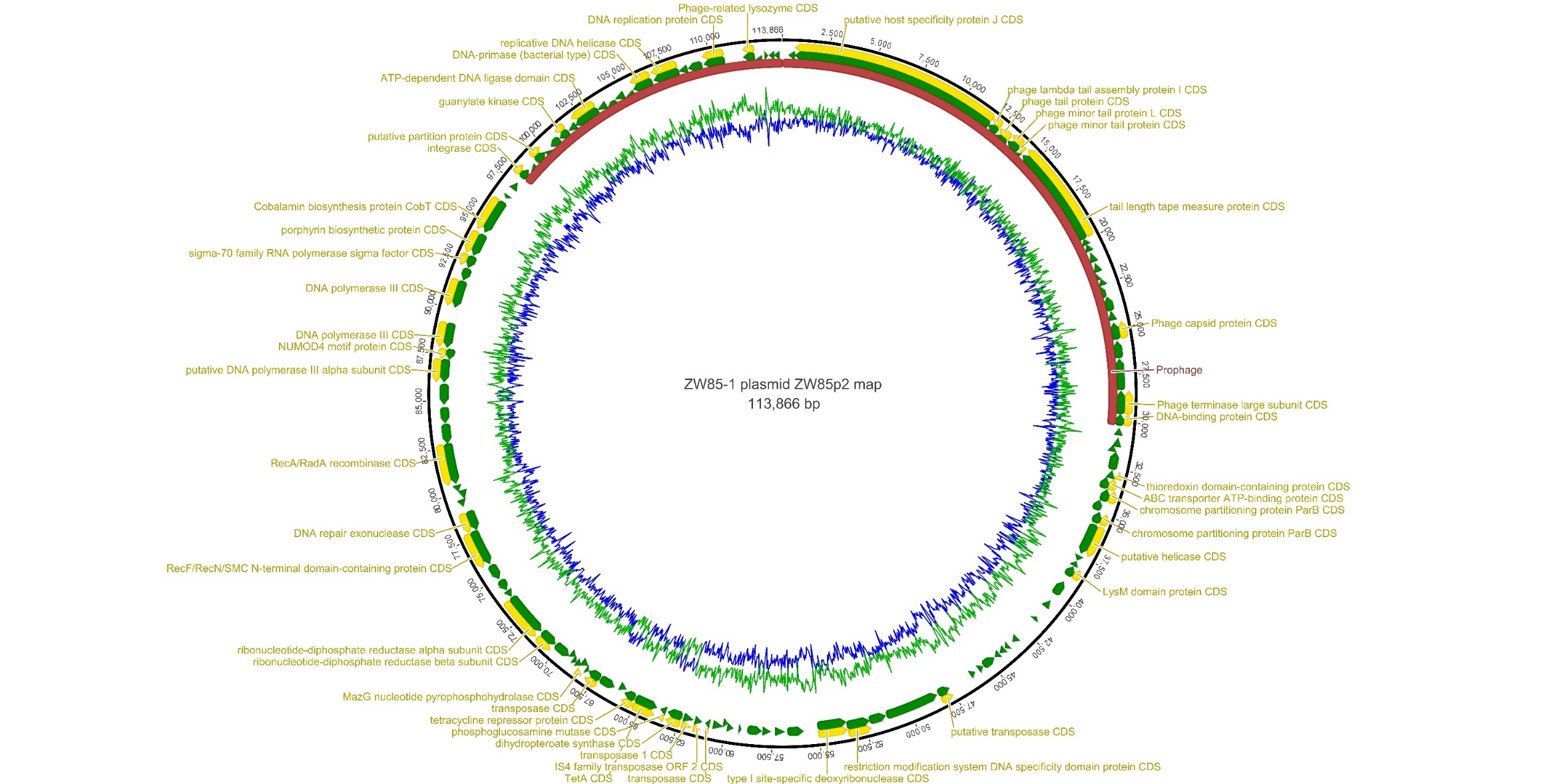

**Figure S1D.** Map of plasmid ZW85p2 of *Acinetobacter baumannii* strain ZW85-1. Green arrows: identified open reading frames; Yellow arrows: annotated coding DNA sequences (CDS); Pink arrow: tRNA; Red region: prophage sequence identified. Graphic for GC (blue) and AT (green) content in center.
