## Supplementary Materials for "Genomic analysis of *Acinetobacter baumannii* prophages reveals remarkable diversity and suggests profound impact on bacterial virulence and fitness"

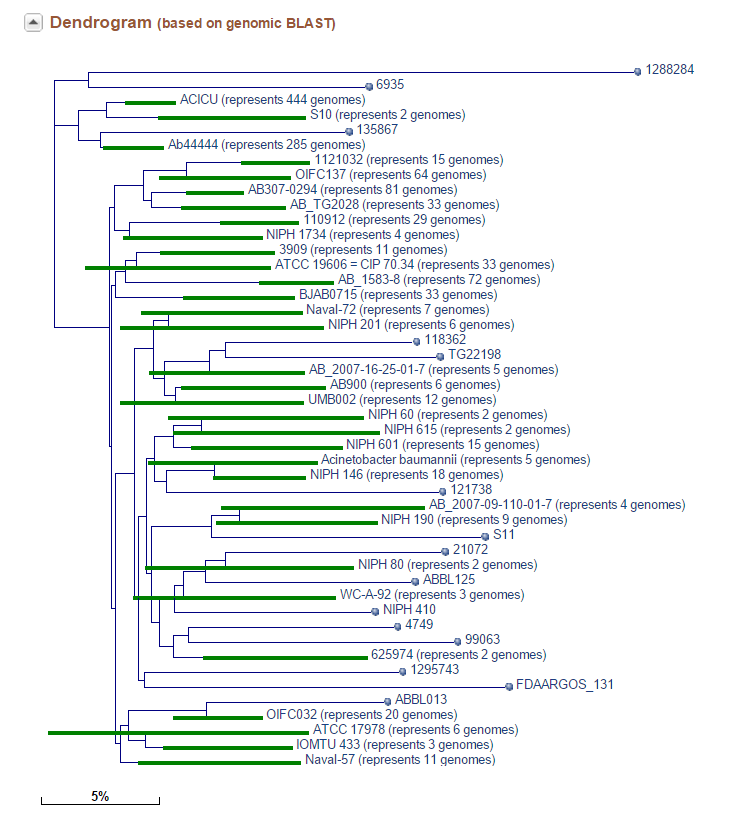

**Figure S2.** Dendogram of *Acinetobacter baumannii* retrieved from GenBank at the date of April 11, 2016. Representative strains from each branch were selected for detailed analysis of encoded prophages.
